# The elite common bean *Phaseolus vulgaris* cultivar Pinto Saltillo hosts a rich and diverse array of plant-growth promoting bacteria in its rhizosphere

**DOI:** 10.1101/2023.10.23.563606

**Authors:** Griselda López-Romo, Rosa Isela Santamaría, Patricia Bustos, Francisco Echavarría, Luis Roberto Reveles-Torres, Víctor González

**Author notes:** Address correspondence to: Víctor González,. Rosa Isela Santamaría and Patricia Bustos contribute equally to this work.

## Abstract

The rhizosphere of crop plants is a nutrient-rich niche that is inhabited by many microorganisms. Root-associated microorganisms play a crucial role in crop yields in agriculture. Given the ample diversity of varieties and cultivars of the common bean (*Phaseolus vulgaris*) used in agriculture, it is important to characterize their bacterial communities. In this study, we analyzed the bacterial rhizosphere components of the bean cultivar Pinto Saltillo, which is widely produced and consumed in Mexico.

Bulk soil and rhizosphere samples from the *P. vulgaris* cultivar Pinto Saltillo were collected *in situ* from plots with and without cultivation history. Metagenomic analysis revealed that in both plots, the bacterial diversity in the bulk soil exceeded that in the rhizosphere. Moreover, diversity and taxonomic composition analysis confirmed the dominance of Proteobacteria in the rhizosphere. Comparisons with pairs of bulk soil-rhizosphere metagenomes of other cultivated plants (maize, wheat, tomato, cucumber, and the model plant *Arabidopsis*) indicated a pronounced rhizosphere effect of the cultivar Pinto Saltillo, particularly regarding the presence of bacterial genera already known as plant growth promoters, including *Rhizobium*. Metagenome-assembled genomes (MAGs) reconstructed from metagenomes confirmed a diverse set of species at the OTU level, closely related to this group of microorganisms. Our analysis underscores the association of *R. sophoriradicis* strains as the primary nodulating agent of common beans in the sampled agricultural fields.

These findings imply that the success of common bean crops relies on microbial species that are still inadequately characterized beyond the established role of nitrogen-fixing bacteria.

**Importance:** Sustainable agriculture is a long-term goal aimed at mitigating the impact of modern intensive and polluting agricultural technologies. Significant efforts are underway to understand the contributions of microorganisms to the health and productivity of crop plants. The common bean (*Phaseolus vulgaris*) is a domesticated leguminous plant native to Mesoamerica, that whose seeds provide sustenance for millions of people in America and Africa. Previous studies have illuminated the bacterial diversity of the rhizosphere microbiome in relation to plant resistance to pathogens and in the domestication process. These findings underscore the importance of investigating the bacterial rhizosphere communities in successful cultivars of the common bean. In this study, we demonstrate that the common bean cultivar Pinto Saltillo hosts a diverse array of plant-growth promoting bacteria in its rhizosphere. These findings suggest that the agricultural success of common bean cultivars could be attributed to the interplay between the plant and its rhizosphere bacterial community, rather than solely relying on nitrogen-fixing symbiosis.

## Introduction

The rhizosphere is the soil region surrounding plant roots (1). It is a nutrient-rich niche inhabited by a large and diverse microbial community, known as the microbiome (2). The rhizosphere microbiome is comprised of many species of bacteria, archaea, fungi, nematodes, protozoa, and viruses. These microorganisms are involved in various ecological interactions and their physiological activities are essential for plant nutrition, development, and pathogen defense (3, 4).

Earlier studies on the rhizosphere have primarily focused on culturing specific bacteria and microorganisms to characterize their contributions to plant growth or to identify their pathogenic behavior. However, metagenomic methods have revealed that the abundance and diversity of rhizosphere microorganisms are more significant than previously thought (5). Consequently, the contribution of the rhizosphere to plant development surpasses that of individual species (5) .

Previous studies on model plants, such as *Arabidopsis* and *Lotus*, as well as agriculturally significant plants, such as rice, maize, wheat, and tomato, have established that the rhizosphere microbiome assembles from the soil under the influence of plant root exudates and rhizodeposition (6–10). Simultaneously, the richness and diversity of microorganisms are typically high in bulk soil, but decrease in the presence of cultivated plants (3, 11, 12). Thus, the interplay between plants and the bacterial community in the bulk soil determines the composition of the rhizosphere microbiome and other compartments, such as the closest microbial community adhered to the plant root at the rhizoplane and the endophyte community in the endosphere (13).

The symbiosis between *Rhizobium* and diverse leguminous plants (Family Fabaceae) is a notable example of the intimate relationship between plant roots and bacteria (14). This symbiosis significantly contributes to nitrogen incorporation into the biosphere by forming N-fixing nodules in the roots. This symbiotic association partially explains the evolutionary success of the Fabaceae family and its use in human alimentation. The domestication of several Fabaceae species dates to at least 10,000 years and continues with new inbreeding techniques (11, 15, 16). Thus, selected genetic features of cultivated plants, such as root architecture and pathogen resistance, likely play an indirect role in favoring the adaptation of specific microbial species to the rhizosphere (17, 18). The rhizosphere microbiome of leguminous plants, such as soybean, clover, and common bean, has been studied only in a few instances (19–21). There is experimental evidence of a reduction in the diversity of the bacterial community from wild type to cultivated common beans (21, 22).

Additionally, the response of the bacterial rhizosphere community across various common bean genotypes and agricultural soils suggests high variability in the composition of the bacterial community and a reduced common and persistent set of genera (23). The taxonomic composition of the bacterial rhizosphere community indicates species dominance from the phyla Proteobacteria, Actinobacteria, and Bacteroidetes (22). Remarkably, contrasting soil types influenced the rhizosphere community, as revealed by the differential composition and abundance detected at lower taxonomic ranks, such as families and genera (22). However, high-resolution studies to identify the composition of the rhizosphere microbial community at the species-OTU level are still lacking.

This study aimed to identify the bacterial rhizosphere community of a single common bean cultivar (*Phaseolus vulgaris* cultivar Pinto Saltillo) that is widely produced and consumed in northern Mexico (24, 25). We hypothesized that the bacterial rhizosphere community of the Pinto Saltillo cultivar played an essential role in the success of this crop. Building on previous findings, we obtained soil and rhizosphere samples directly from agricultural plots with and without prior cultivation history. We used shotgun metagenomic methods to thoroughly explore bacterial diversity at the species-level OTUs.

Our study revealed a strong influence of the Pinto Saltillo cultivar on the composition of the bacterial rhizosphere community, regardless of the soil type and cultivation history at agricultural sites. The community is predominantly composed of species from the phylum Proteobacteria, including several Plant-Growth Promoter Rhizobacteria (PGPR), which may be an essential element for the agricultural success of the Pinto Saltillo cultivar.

## Materials and Methods

### Sampling Description

This study was conducted in the experimental field of the Instituto Nacional de Investigaciones Forestales, Agrícolas and Pecuarias (INIFAP) in Zacatecas, Zacatecas, Mexico. Two adjacent agricultural plots, one referred to as “N” for “non-agricultural,” which had no history of cultivation but had been used for tillage tests with agricultural machinery for several years [22°54’20.932 “N 102°39’29.504 “W]. The second plot, denoted as “A” or “agricultural,” had a history of cultivation with a crop rotation system (corn, beans, and chili) for an extended period [22°54’23.8 “N 102°39’30.4 “W] (Supplementary Figure S1). The total area of the plots was 0.34 and 0.42 hectares for sites N and A, respectively. Both sites were planted with the common bean cultivar “Pinto Saltillo” in July 2020.

In each plot (“N” and “A”), we established a grid of 6 × 12 meters, which was further divided into 18 quadrants of 2 × 2 meters (Supplementary Figure S1). Bulk soil samples were collected from ten these quadrants for physicochemical analysis. Bulk soil samples (before planting) and rhizosphere samples post-planting were obtained from ten quadrant intersections labeled with letters A to J in both plots, ensuring that the same points in bulk soils corresponded to the locations where bean seeds were sown. Rhizosphere soil was collected after 40-45 days post-planting at the flowering stage. For the metagenomic experiments, bulk soil (stage 1) and rhizosphere (stage 2) samples were collected from nine plants. All samples were stored at -20°C until DNA extraction, but only 17 samples were processed for DNA purification and subsequent metagenomic experiments (Supplementary Table S1). Bulk soil samples were collected using metal core augers (10 cm length and 4.3 cm diameter) at a depth of 10 cm. Twenty samples were collected per plot, ten of which were used to determine the physicochemical characteristics of the soil.

In this study, the rhizosphere was defined as the soil firmly attached to the root (7). Therefore, we removed the soil surrounding the roots, leaving only the tightly adhered soil. Then, the roots were shaken vigorously in a bag, transferred to a conical tube with SM buffer (100 mM NaCl, eight mM MgSO4 · 7H2O, and 50 mM Tris-Cl, pH 7.5), and shaken vigorously again with a vortex. Subsequently, the roots were removed, the tubes were centrifuged for 10 min at 10,000 rpm, and the supernatants were discarded. The soil pellets were used for DNA purification.

### DNA Purification

The soil or rhizosphere (250 mg) was used for DNA extraction using the DNeasy PowerSoil DNA Isolation Kit (Qiagen), following the manufacturer’s protocol, and stored at -20°C until analysis. Seventeen samples were sequenced: three bulk soil samples from plot A, three bulk soil samples from plot N, six rhizosphere samples from plot A, and five rhizosphere samples from plot N (Supplementary Table S1). All samples were sent to Macrogen for Shotgun sequencing using the TruSeq kit on Illumina NovaSeq, with a sequence length of 150 bp paired-end reads.

### Physicochemical Characteristics of the Soil

Soil samples from both plots were classified as silty clay loam, clay loam, silty clay, and clay, which corresponds to the commonly described classification for this soil type (22, 26). Chemical characteristics related to soil fertility of the eleven meta-samples were classified as medium alkaline (pH 8-8.49), with a medium proportion of organic matter (1.39 - 3.65 %), low inorganic nitrogen (4.67 - 20.97 mg/kg), and with a high level of phosphorus determined with the Olsen method in the case of the agricultural plots (15.98 - 20.47 mg/kg) and a low to medium level in the non-agricultural plots (15.98 - 24.47 mg/kg). The chemical and physical properties are described in Supplementary Table S2, and the differences between the A and N sites, determined by PCoA statistics, are presented in Figure 1A.

**Figure 1.**
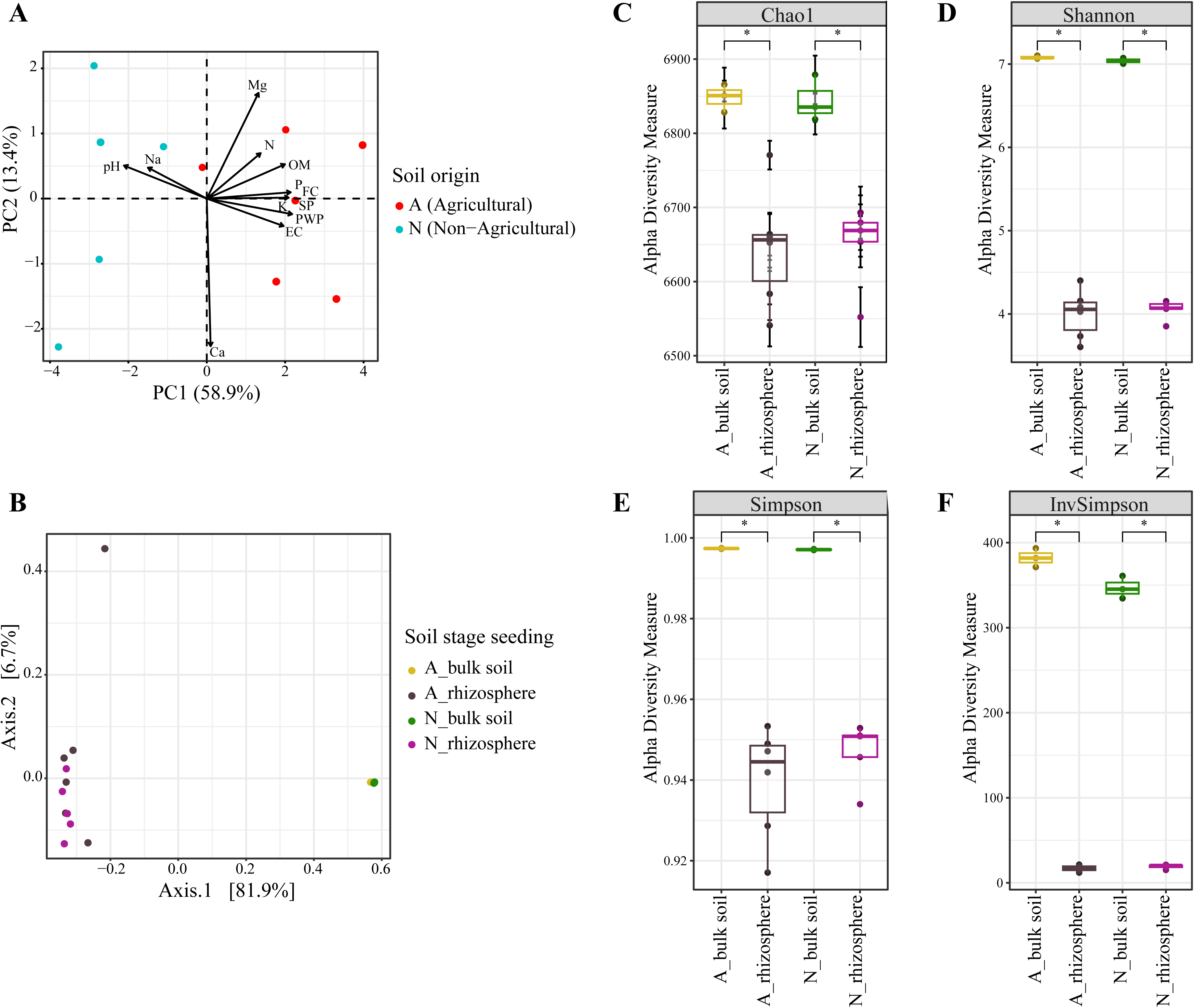
Bacterial diversity in soil and rhizosphere. (A) Principal Component Analysis (PCA) of the soil chemical (pH, N, Mg, K, and P) and physical (OM, FC, SP, PWP, and EC) properties. (B) Principal Coordinate Analysis (PCoA) of beta-diversity based on Bray-Curtis dissimilarity between bacterial populations from the soil and rhizosphere of *P. vulgaris* cultivar Pinto Saltillo (PERMANOVA, P= 0.001). Alpha diversity is shown in panels (C) Chao1, (D) Shannon H’, (E) Simpson D, and (F) Inverse Simpson, 1/D indices. Wilcoxon test p=0.024 in plot A and p=0.036 in plot N are indicated by asterisks.

### Metagenomic Sequencing and Taxonomic Assignment

An average of 99,153,080 and 81,811,594 sequence reads were obtained for soil and rhizosphere samples, respectively (Supplementary Table S3). Sequences with quality scores (Q) greater than 20 were processed using FastQC v0.11.8 (27) and TrimGalore v0.6.4R (28). High-quality reads were taxonomically classified using the Kraken2 program with default parameters: confidence 0.0, minimum-hit-groups 2 (29), and PlusPFP database (Standard plus protozoa, fungi and plant, fixed from 12/2/20 version). Rarefaction curves without replacement were generated using the Vegan package in R, with a *recurve* function (30). The Phyloseq package was employed to estimate alpha diversity (utilizing the estimate_richness function) based on the number of observed taxa, Chao1, and the Shannon and Simpson indices. Beta diversity was assessed using the Bray-Curtis distance method (utilizing distance and ordinate functions) (31). Differential abundance estimates were calculated using the DESeq2 program (32) and visualized using the EnhancedVolcano package in R (33) (Supplementary Table S4).

Reference metagenomes for *Arabidopsis* (NCBI BioProject number PRJNA435676; (34), common bean (PRJNA904562; (35), maize (PRJNA678469 and PRJNA678475; (36), cucumber, wheat (PRJNA208116; (37), and tomato (PRJNA603603; (10) were downloaded from the RSA site in GenBank. The accession numbers of SRA are provided in Supplementary Table S5. The reference metagenomes were processed as described above, including the diversity analysis.

### Metagenome-assembled genomes (MAGs)

High-quality sequence readings from 22 metagenomic sequencing experiments (six from bulk soil, 11 from rhizosphere, and five from plant nodules) were assembled using SPAdes v3.13 and MEGAHit v1.2.8 (38, 39). The assemblies produced by both programs were compared with the QUAST v5.0.2 program (40), selecting the assemblies with the largest N50 and contigs. For bulk soils and rhizosphere, the best assembler was MEGAHIT, whereas for nodule samples, the best assembler was SPAdes. Contig coverage was obtained by aligning high-quality reads to contigs using Bowtie2. Contig binning was performed using CONCOCT 1.1.0 (41, 42), MaxBin2 2.2.7 (43), MetaBAT1 1.2.15 (44), MetaBAT 2.2.15 (45), and their results were incorporated into the ANVIO’s (46), suite to evaluate completeness and redundancy. Redundancy in the MAGs prediction among the binning programs was solved by selecting the largest with at least 60% completeness and 10% redundancy. A total of 150 MAGS were selected:113 from the rhizosphere, 21 from the nodules, and 16 from the bulk soil (Supplementary Table S6). They were taxonomically assigned using the Genome Taxonomy Database Toolkit GTDB-Tk 2.1.1 (47). *Rhizobium* MAGs were compared to the reference strains CFN42, CIAT652, Kim5, and CCBAU 03470 using the (*averaverage_nucleotide_identity. py* of pyani (48).

### Statistics

Permutational multivariate analysis of variance (PERMANOVA) was performed with 999 permutations using the Phyloseq package with distance function and adonis2 function from the Vegan package (30, 31). Wilcoxon and t-student were made with Stats package (49). In the case of the PCoA analysis were made with the ordinate function from the Phyloseq package (31).

### Data availability

Raw sequence data are available at the National Center for Biotechnology Information in the Sequence Read Archive (SRA) section under the accession number PRJNA1015279.

## Results

### Site sampling and physicochemical composition analyses

To compare the bacterial diversity associated with the soil and rhizosphere of common bean (*P. vulgaris* cv. Pinto Saltillo), samples were collected from plots with agricultural history over the past five years (site A) and from plots without prior agricultural use (site N), adjacent to site A (Supplementary Figure S1). Ten soil samples from each site were analyzed for their physical characteristics and chemical composition, revealing significant differences within and between sites (Figure 1A; Supplementary Table S2). Key chemical contributors to inter-site differences included factors such as pH, nitrogen, and phosphorus, while physical attributes such as moisture (SP, Saturation Point %), PWP (permanent-ilting-point %), and FC (Field Capacity %) also displayed variations within the sites (Figure 1A).

### High-resolution shotgun sequencing

Our study aimed to achieve the highest representation of the bacterial community in the bulk soil and rhizosphere of Pinto Saltillo bean. Consequently, to ensure the most comprehensive coverage of the resident species in such communities, we obtained an average of 87 Gb of shotgun paired-end sequences per bulk soil sample (three per site, N, and A) and rhizosphere (six for site A and five for site N). Overall, the GC profile in the bulk soil was higher (65%) than in the rhizosphere (60%), suggesting that they were composed of contrasting bacterial communities.

Sequence reads were classified into OTUs using Kraken2 with default settings to maximize the recall (29). Even so, in the bulk samples, the percentage of classified reads relative to unclassified sequence reads was low (20%), whereas in the rhizosphere, the classified reads reached an average of 78% (Supplementary Table S3).

The bacterial domain exhibited the highest percentage of classified sequence readings at sites A and N, either bulk soil or the rhizosphere (98%). The remainder of the sequences belonged to archaea, viruses, fungi, and other eukaryotes (see Supplementary Table S3). In all metagenomic samples, the breadth of bacterial OTUs reached a plateau in rarefaction statistics, indicating sufficient sampling depth and coverage to estimate the diversity (Supplementary Figure S2).

### Bacterial diversity in soil and rhizosphere

To obtain evidence regarding the taxonomic structure and relative abundance of the bulk soil and rhizosphere bacterial communities, we employed three diversity indices using PhyloSeq (Chao, Shannon, and Simpson; see (31)). In the bulk soil at site A, the median diversity indices of the three samples were Chao1 = 6,851 OTUs and Shannon H’ = 7.07. Site N did not exhibit substantial differences in diversity compared to site A (Chao1 = 6,835 OTUs and Shannon H’ = 7.04; Wilcoxon p = 0.2) (Figure 1C). Furthermore, although the Chao1 values were similar (6,656 and 6,669 OTUs, respectively) for the bacterial communities in the rhizosphere of sites A and N, the diversity measured by the Shannon index was notably lower (H’ = 4.05 for site A and 4.07 for site N) than for the bulk soil at the two sites (Figure 1D). Therefore, the results suggested that the diversity of the bacterial community in the soil before planting surpassed that found in the rhizosphere (Wilcoxon p = 0.024 and 0.036 for A and N within-site comparisons).

Moreover, we calculated the Simpson index D and its inverse (1/D) for all the samples (Figure 1E-F). The median D of the soil samples was 0.997 (1-D = 0), whereas in the rhizosphere, the values had a range of D = 0.917-0.953 (median D = 0.947) in both plots (A and N). Accordingly, at sites A and N, the inverse Simpson index of the bacterial community in the bulk soil ranged from 371 to 393 and from 334 to 360 (medians of 382 and 345 for A and N, respectively). In contrast, the inverse Simpson index of the rhizosphere samples was significantly lower (12.05 to 21.43 and 15.15 to 21.22 for A and N sites, with a median of 18 and 20). Thus, the bacterial community was more homogeneous in the soil than in the rhizosphere, but dominance of some OTUs could be expected in the rhizosphere.

Considering that the soil and rhizosphere compartments may harbor different bacterial communities, we aimed to compare them and quantitatively assess their differences. We then calculated beta diversity based on the Bray-Curtis dissimilarity between OTUs. Principal coordinate analysis (PCoA) of beta diversity showed that soil and rhizosphere bacterial communities were distinctly different and did not overlap. Differences in beta diversity between these communities explained 81.9% of the variance in the PCoA (PERMANOVA analysis, P < 0.001; Figure 1C). The bacterial rhizosphere communities of sites A and N did not display significant differences from each other, except for the sample labeled AH, which was significantly different from the rest of the samples.

### Bacterial taxonomic compositions in the soil and rhizosphere

To assess the taxonomic composition of the soil and rhizosphere bacterial communities, taxon ranks were assigned to metagenomic reads using Kraken2, a free-alignment algorithm based on k-mers (29). Bulk soil samples from sites A and N had similar OTU compositions. In the bulk soil, the phyla Actinobacteria and Proteobacteria accounted for the highest percentage of 54.37% and 58.78% of classified readings, and 40.41 % and 35.31% for A and N sites, respectively (Supplementary Figure S3). Members of the phyla Firmicutes, Planctomycetes, and Bacteroidetes were less represented (2.89% and 3.10% for the A and N sites, respectively). Among the most abundant genera in the soils were *Streptomyces* (9.91%), *Nocardiodes* (4.95%), *Micronospora* (2.78%) of phylum Actinobacteria; and *Sphingomonas* (2.74%), *Bradyrhizobium* (2.48%), and *Pseudomonas* (1.77%), of the Proteobacteria (Figure 2).

**Figure 2.**
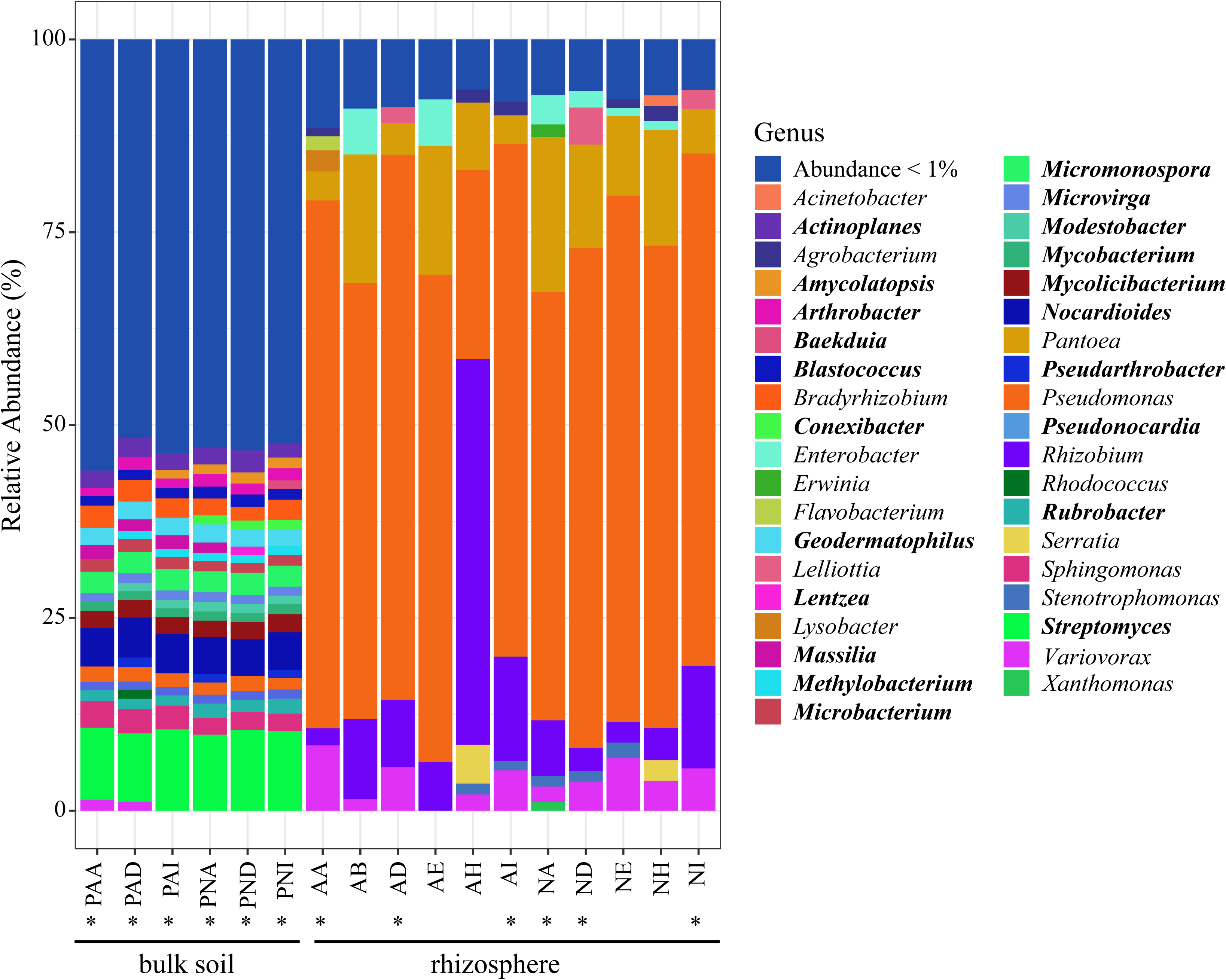
Bacterial taxonomic composition and relative abundance of the bulk soil and bean rhizosphere at the genus level. Stacked bars represent the Kraken2 taxonomic classification and abundance of sequence reads of metagenomic samples from bulk soil and the rhizosphere from agriculture (A) and non-agricultural (N) sites. The nomenclature for each metagenome used a three-letter system for bulk soil samples taken before planting (P) at sites (A) and (N). The last letter indicates the intersection coordinates of the grid (Supplementary Figure S1). Samples from the rhizosphere were identified by two letters pointing to sites A and N, and the coordinates in the grid. Metagenomic samples from the bulk soil corresponded exactly to sites where the rhizosphere samples were subsequently taken from the plant roots (indicated by asterisks). The right inset shows the colors according to genera with abundance > 1%; low-represented genera (< 1%) are shown together. Genus names in bold correspond exclusively to bulk soil.

Proteobacteria dominated the rhizosphere community (98.22%). A minor proportion of the reads was assigned to Actinobacteria (1.02%) and Bacteroidetes (0.57%) (Supplementary Figure S3). When the proportions of the taxonomic categories within Proteobacteria were evaluated, the class gamma-Proteobacteria (79.24%) and the genus *Pseudomonas* dominated the read assignments, and in less proportion, the alpha- and beta-Proteobacteria (13.78% and 5.11 %, respectively) (Supplementary Figure S4). The genus *Pseudomonas* appeared to dominate the rhizosphere community, accounting for the highest proportion (60.69%). A set of six other genera that represented > 1% of the sequence reads, including *Rhizobium* (11%, Alpha-proteobacteria), *Pantoea (*10%, Gamma-proteobacteria), *Variovorax* (4%, beta-proteobacteria), *Enterobacter* (2%, gamma-proteobacteria), *Serratia* (1%, gamma-proteobacteria), and *Lelliotia* (1%, Gamma-proteobacteria), accounted for 30% of the assigned reads (Figure 2). The remaining phyla and genera were present in low proportions (< 1%) and were composed of some genera of alpha-proteobacteria and unknown bacterial taxa. Differences in the bacterial composition of the soil and rhizosphere communities were apparent across all taxonomic hierarchies.

### Shifts in OTUs composition from bulk soil to rhizosphere

To quantify the differences between the bacterial communities in the bulk soil and rhizosphere, we conducted a differential abundance comparison using DESeq2 (Figure 3). There were 4,856 (69%) OTUs showing changes in their relative abundance upon the introduction of the common bean plant (Figure 3A; Supplementary Table S4). Almost half of these OTUs (2,604) increased their representation in the rhizosphere community compared to that in the bulk soil. Notably, the majority of the OTUs (1,922, 39.6%) belonged to the phylum Proteobacteria, with smaller proportions in Firmicutes (209, 4.30%) and Bacteroidetes (395, 8.13%). OTUs of Proteobacteria were taxonomically assigned to 321 genera, whereas Firmicutes and Bacteroidetes were included in 70 and 113 genera, respectively.

**Figure 3.**
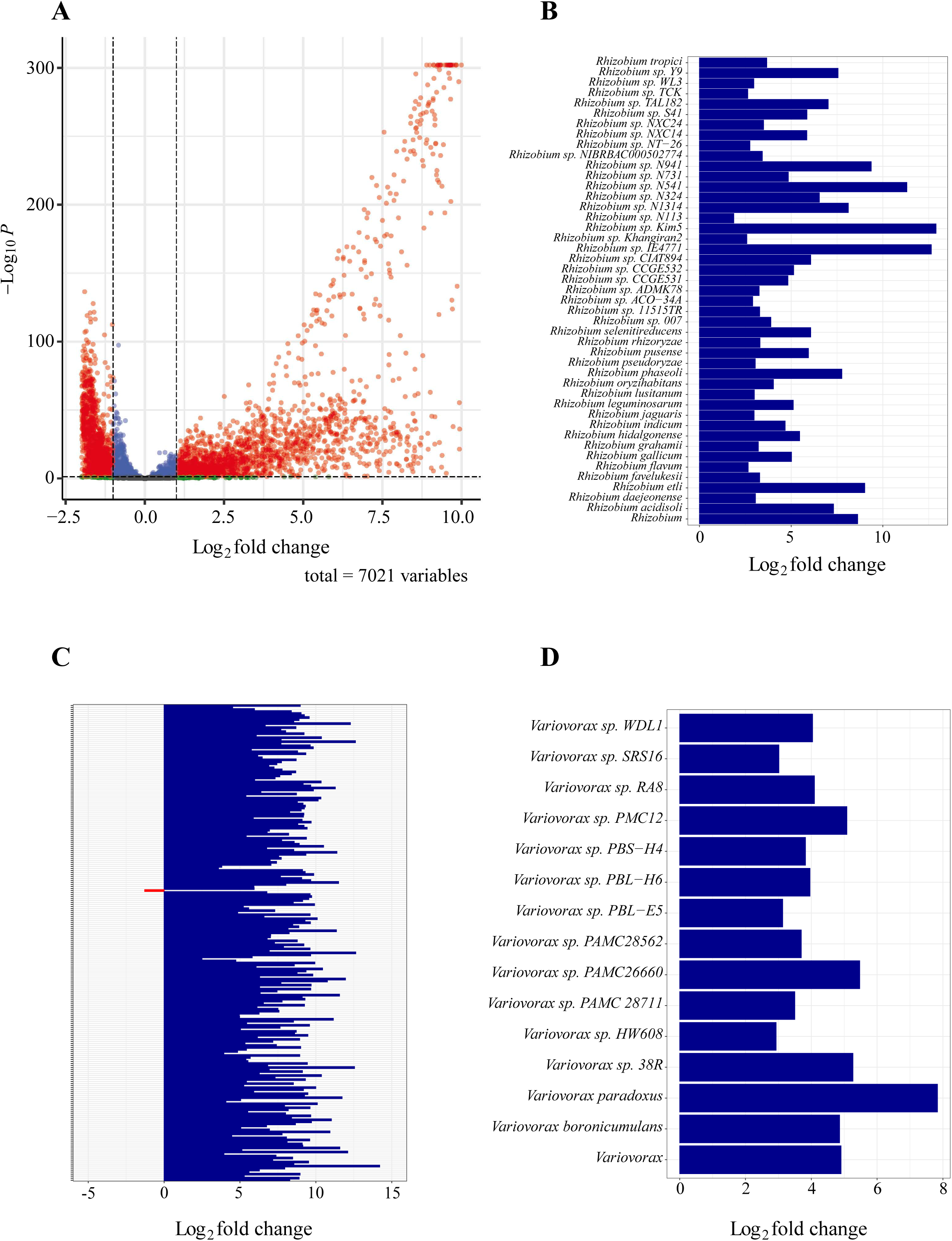
Shifts in bacterial composition from the bulk soil to the rhizosphere. The differential abundance of bulk soil and rhizosphere OTUs was evaluated using DESeq2. (A) Volcano plot of differential abundance expressed as log2 (fold change, *x*-axis) and its statistical significance p-value (-log10, *y*-axis). A total of 7,021 OTUs were tested for differential abundance, but only 4,856 showed significant changes (p < 0.05), as indicated by red dots. Parallel dashed lines delimited the area containing less significant changes (p > 0.05, log2-fold < 1). Log2-fold change for the OTUs assigned within the genera *Rhizobium* (B), *Pseudomonas* (C), and *Variovorax* (D)—the species names for the OTUs were those assigned by Kraken2. Species names are omitted in (C) because of the ample number (237) in *Pseudomonas*. In red, OTUs with significance (log2-fold > one and a p-value < 0.05); in blue are those non-significant OTUs that have a p-value < 0.05, but log2-fold < 1; in green are those non-significant with Log2-fold > 1 but p-value > 0.05, and in gray are the non-significant OTUs that do not meet either of the two parameters (log2-fold < 1 and p-value > 0.05).

Remarkably, OTUs related to plant-growth-promoting rhizobacteria (PGPR) proliferated in the rhizosphere. Specifically, within Proteobacteria, OTUs identified within the genera *Pantoea, Lelliottia, Enterobacter, Erwinia, Pseudomonas, Serratia, Acinetobacter,* and *Scandinavium*, all parts of the gamma-proteobacteria class, showed an abundance increase ranging from 7-to 14-fold. Among the alpha-Proteobacteria populations, genera with significant Log2 fold changes included *Rhizobium, Agrobacterium, Neorhizobium, Sinorhizobium,* and *Ensifer*, all enriched 3-to 12-fold. Additionally, the genera *Variovorax* and *Achromobacter*, belonging to gamma-proteobacteria, displayed seven and 4-fold increases, respectively. Only a few OTUs affiliated within a single bacterial genus significantly increased their proportion in the rhizosphere (Figure 3B-D). In *Rhizobium*, only three out of 46 OTUs assigned by Kraken2 accounted for the highest sequence read abundance in the rhizosphere (Supplementary Figure S5). This was also shown by the increased log 2-fold change from three to ten in every *Rhizobium* OTU (Figure 3B). The same pattern was observed in other rhizosphere bacteria such as *Pseudomonas* and *Variovorax* (Fig. 3C-D)

In contrast, 1,169 OTUs (24.07%) belonging to Actinobacteria species in the bulk soil exhibited significant decreases in the rhizosphere community. Other OTUs from Proteobacteria (10.36%), Firmicutes (3.67%), Cyanobacteria (2.24%), and Bacteroidetes (1.17%), represented by 503, 178, 109, and 57 taxa, respectively, also underwent reductions in their presence in the rhizosphere (Figure 3).

Altogether, these results indicate that common bean plants significantly influence the rhizosphere community by stimulating the growth of a specific array of bacterial species.

### Rhizosphere effects on other plants

We compared the responses of the bulk soil and the rhizosphere of other cultivated plants to elucidate the specific impact of the bean cultivar Pinto Saltillo on the rhizosphere bacterial communities. To achieve this, we utilized metagenomic reads available in the NCBI-GenBank for several plants, including the common bean (*P. vulgaris* cultivars IAC Milenio and IAC Alvorada), the model plant *Arabidopsis thaliana* (Brassicaceae), maize (Poaceae, *Zea mays*), wheat (Poaceae, *Triticum turgidum* cv. Negev), tomato (Solanaceae; *Solanum lycopersicum* L. cv. Río grande), and cucumber (Cucurbitaceae; *Cucumis sativus* cv. Kfir-413). The analysis of the diversity and taxonomic composition of this dataset revealed that in these plants, species of Proteobacteria cohabit the rhizosphere niche with Actinobacteria species (Supplementary Figure S6). Furthermore, differences were observed in the bacterial composition of the rhizosphere of these plants at lower taxonomic levels, such as class and genus. In the rhizosphere of the common bean Pinto Saltillo, the class gamma-Proteobacteria dominated the community, whereas in the rhizospheres of other plants, alpha- and beta-Proteobacteria, along with some classes of Actinobacteria, were more abundant in the rhizosphere communities (Supplementary Figure S8). Notably, only a subset of bacterial genera exhibited shifts in abundance from the soil to the rhizosphere of the reference plants, excluding *Pseudomonas*, *Rhizobium*, and other genera that thrive in the common bean Pinto Saltillo. Although OTUs belonging to *Rhizobium* were present in all seven rhizosphere communities, substantial shifts in the abundance of *Rhizobium* from bulk soil to the rhizosphere were only observed in the common bean rhizosphere but were moderate in non-leguminous plants (see Supplementary Figure S8) (Figure 4).

**Figure 4.**
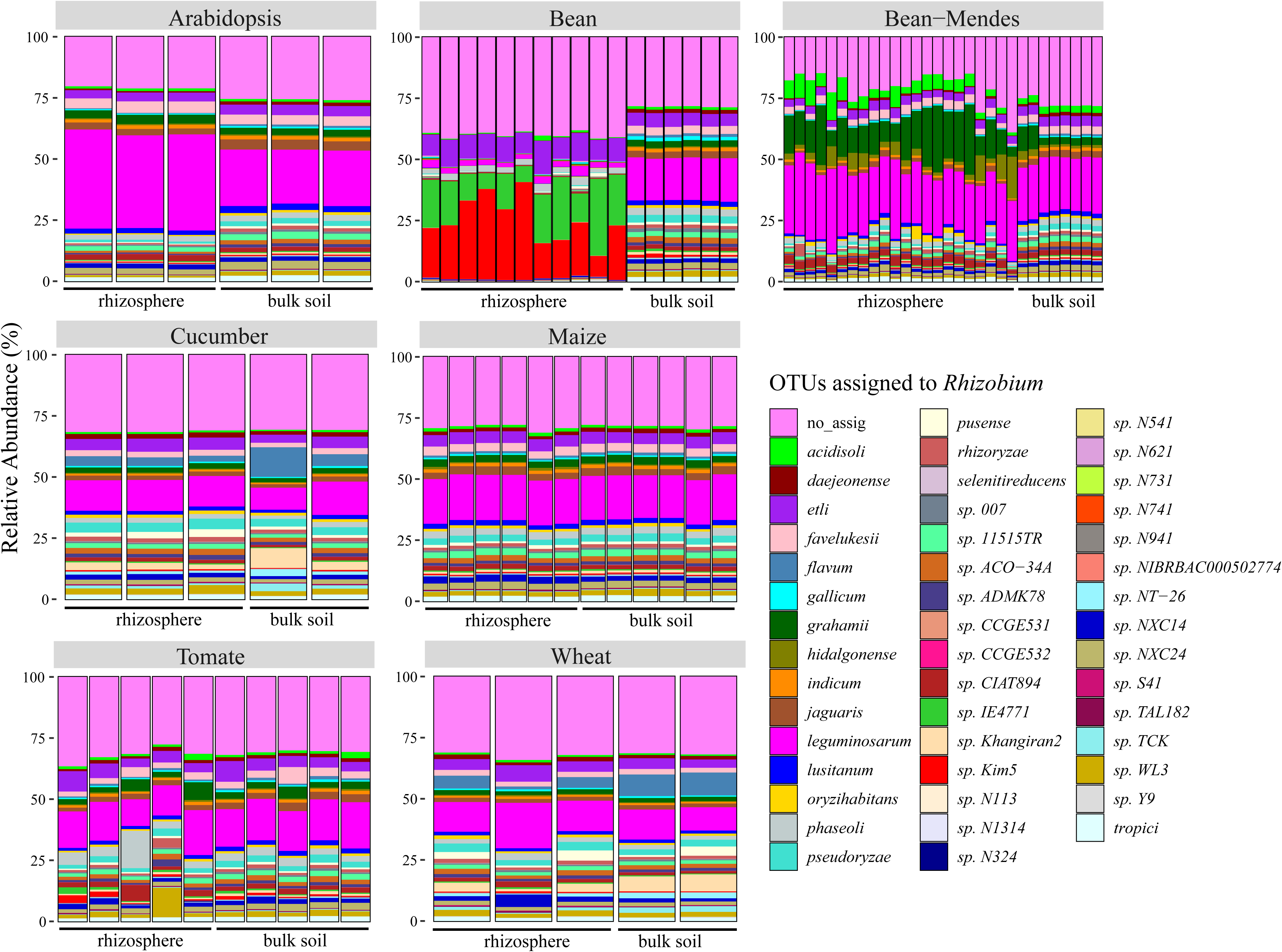
The rhizosphere effect in common beans and other plants is focused on the *Rhizobium* genus. Pairs of bulk soil and rhizosphere metagenomes of six cultivated plants were downloaded from the RSA archive of GenBank and used to compare the enrichment of *Rhizobium* OTUS assigned with Kraken2. Stacked bars show abundance relative to the rhizosphere and bulk compartments of *Arabidopsis thaliana* (Stringlis et al. 2018), common bean (*Phaseolus vulgaris* cv. Pinto Saltillo), common bean (*Phaseolus vulgaris* cv. IAC Milenio and cv. IAC Alvorada; Mendes et al., 2023), cucumber, and wheat (*Cucumis sativus* cv. Kfir-413 and *Triticum turgidum* cv. Negev, (37), maize (*Zea Mays*, (36) Babalola et al. 2021), and tomato (*Solanum lycopersicum* L. cv. Río Grande, (10). The inset shows the taxonomic assignations to putative *Rhizobium* species.

### Bacterial Community Structure

Bacterial species within microbial communities comprise diverse populations of closely related but not identical individuals. We assessed the structure of the bacterial community by reconstructing Metagenome-Assembled Genomes (MAGs) (see Methods). To achieve this, we employed various bioinformatics methods to bin the metagenomic contigs obtained from the soil and rhizosphere of two sites: A and N (see details in Methods). We successfully reconstructed large MAGs meeting the high- and medium-quality criteria (>1 Mb, completion >60%, and redundancy <10%), as depicted in Supplementary Figure S9. A total of 150 high-quality MAGs were obtained from all the metagenomic samples (Figure 5, Supplementary Table S6). Among these, 16 MAGs were isolated from bulk soil, representing the agricultural soil community before planting, and 113 were reconstructed from the rhizosphere community. The remaining 21 MAGs were reconstructed from the root nodules. Notably, OTUs assigned by Kraken2, with the most abundant sequence read representation, were often associated with several reconstructed MAGs. However, it is essential to highlight that MAGs comprised of fewer OTUs.

**Figure 5.**
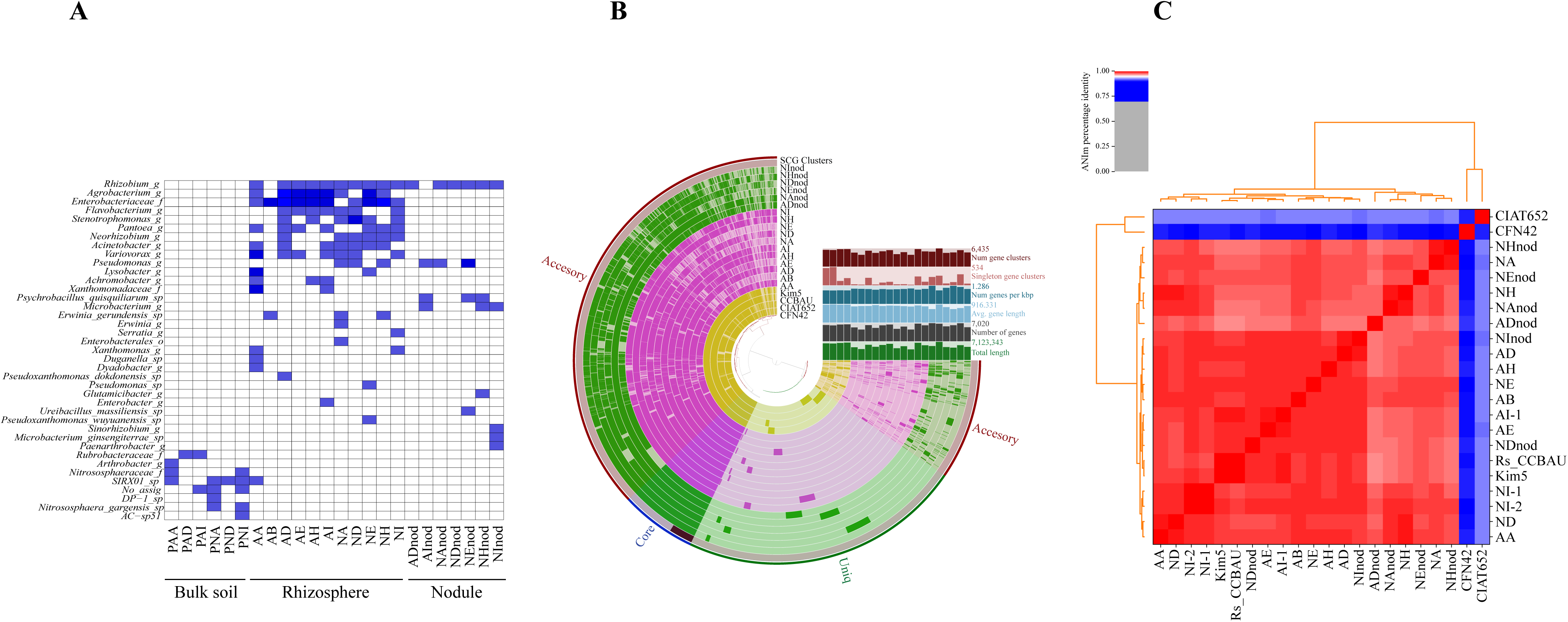
MAGs from bulk soil, rhizosphere, and nodules. (A) Distribution matrix of high- and medium-quality MAGs (completeness > 60%; redundancy < 10%) according to their origin from metagenomic samples of the rhizosphere, bulk soil, and nodule compartments. Blue squares indicate the presence of one MAG, and intense blue squares indicate that two MAGs of the same species were reconstructed from the same metagenomic sample. The GTDB taxonomy assigned to the MAGs is at the left of the matrix. The nomenclature of the metagenomic samples is shown in Figure 2. Metagenomic samples from nodules are indicated with the “nod” word after the name of the metagenome of origin. (B) Meta-pangenome of *Rhizobium* represented by the ANVI’O’s plot. The gene content of *Rhizobium* complete genomes was compared with that *of Rhizobium MAGs using an anvi-pan-genome* (DIAMOND & MCL) within the ANVI’O suite. In concentric circles, from the center to the outermost circle, the whole gene content in the complete genomes of known *Rhizobium* species (yellow circles), MAGs genes from the rhizosphere metagenomes (cherry circles), and MAGs from nodule metagenomes (green circles). The outermost circles represent the single-copy gene clusters (SCG). The single black bar in this circle denotes ribosomal genes. The outermost circle corresponds to the meta-pangenome components: core, accessory, and singleton genes. (C) Comparison of the average nucleotide identity by MUMmer (ANIm) comparison of the complete reference genomes of *Rhizobium* strains (CIAT652, CFN42, Kim5, and CCBAU 03470) with *those of Rhizobium* MAGs. The upper inset shows heatmap identity values as percentages.

Before planting, the bulk soil harbored a dominant Actinobacteria community, as mentioned earlier. Accordingly, the MAGs originating from the soil samples were affiliated with Actinobacteria, including the Micrococcacea and Rubrobacteraceae families. Furthermore, MAGs affiliated with ammonia-oxidizing Archaea of the Nitrososphaeraceae family (Thermoproteota phylum) were of particular significance. Its role in the rhizosphere of plants is unknown, but its potential implications for plant health and nutrient cycling warrant further investigation.

A total of 113 MAGs were successfully retrieved from the rhizosphere, encompassing representatives from 14 genera of gamma-Proteobacteria, three from alpha-Proteobacteria, one from Actinobacteria, and one from Bacteroidetes (Figure 5A). Notably, among these MAGs, several were assigned to bacterial genera commonly considered within the Promotor-Growth Plant Rhizobacteria category, including *Rhizobium, Pantoea, Agrobacterium, Variovorax, Flavobacterium, Pseudomonas, Stenotrophomonas,* and *Acinetobacter*. This bacterial collection highlights a possible influence of Pinto Saltillo on crop success in fields in northern Mexico. Exploring the rhizosphere microbiome of Pinto Saltillo in highly contrasting agricultural soils may provide insights into the reliability of this group of bacteria.

### Identification of *Rhizobium* species associated with the Pinto Saltillo cultivar

The identification of *Rhizobium* species associated with the cultivar Pinto Saltillo is crucial. Although bacterial taxonomic identification at the genus level provides a broad overview of the rhizosphere microbiome, it lacks the resolution needed to discern species and strains, which are fundamental for understanding the structure and function of the rhizosphere. For example, there was an uneven distribution of readings assigned to *Rhizobium* OTUs by Kraken2. This distribution followed a sigmoid pattern in the log10 plot (Supplementary Figure S5). Additionally, Log2 fold-change analysis revealed a similar pattern to that of bulk soil OTUs (Figure 3). This indicates that some *Rhizobium* OTUs are enriched in the rhizosphere, whereas others remain underrepresented.

The taxonomic assignment provided by Kraken2 indicates that 46 OTUS were assigned to different *Rhizobium* species. However, overrepresented OTUs (> 100,000 sequence reads) were identified as *R. etli, R. leguminosarum*, and *R. phaseoli* strains. Furthermore, the most abundant *Rhizobium* OTUs were not identified in specific species. Nevertheless, Metagenome-Assembled Genomes (MAGs) corresponding to *Rhizobium* were obtained from all 11 rhizosphere metagenomic samples, providing compelling evidence of their pivotal role in nitrogen-fixing symbiosis with plants. These *Rhizobium* MAGs exhibited sizes ranging from 4,156,564 bp to 6,740,240 bp (median 5,385,442 bp), with a GC content of 61% compared with known *Rhizobium* genomes.

A comprehensive comparison of *Rhizobium* MAGs using the ANVI’O suite with the reference complete genomes of *R. etli* (CFN42), *R. phaseoli* (CIAT652), *R. sophoriadicis* CCBAU 03470, and *Rhizobium* spp. Kim5 revealed a shared core genome characterized by high conservation along with a moderate degree of heterogeneity in the accessory genome, as depicted in Figure 5B. Furthermore, when subjected to ANIm (Average Nucleotide Identity by MUMmer) comparison, *Rhizobium* MAGs demonstrated high identity (>98%) with both *Rhizobium* spp. Kim5, and *R. sophoriradicis* CCBAU 03470, which were previously identified. Moreover, they exhibited substantial differences (>92% and < 96%, respectively) with *R. etli* CFN42 and *R. phaseoli* CIAT652, two widely studied symbionts (50). In agreement with this, metagenome sequencing of the bacterial nodule content from seven common bean plants showed that the *Rhizobium* MAGs assembled from nodule sequences were similar to the rhizosphere MAGs and highly similar to *Rhizobium* spp. Kim5 and *R. sophoriradicis* CCBAU 03470 (Figure 5).

## Discussion

In this study, we characterized the bacterial rhizosphere components of the common bean *Phaseolus vulgaris* cv. Pinto Saltillo. A detailed analysis of the metagenomic sequence read abundance, taxonomy assignments, and MAG reconstruction revealed species-OTU level classifications for several bacterial species. These findings indicate the dominant presence of species in the phylum Proteobacteria. Within Proteobacteria, the PGPR group included *Pseudomonas, Pantoea, and Flavobacterium*, and, as expected, *Rhizobium* was the most differentially abundant species. In-depth sequence analysis revealed that within a genus, specific OTUs were over-represented, despite their wide variability in the community. We concluded that the common bean rhizosphere might select the most adapted OTUs for the niche, which might have significant physiological consequences for the plant.

Agricultural soils are human-adapted ecosystems designed to facilitate the cultivation of domesticated plants. Crop success largely depends on the soil microbiome, which may influence the composition of the plant rhizosphere (51, 52). Therefore, ensuring a precise comparison of the bacterial communities in both the bulk soil and the plant rhizosphere is crucial. In our study, an *in situ* sampling experiment was meticulously planned so that the bulk soil precisely corresponded to the site where the common bean seeds were subsequently sown (see Supplementary Figure S1). Other studies have had distinct considerations for defining bulk soil (22, 23). For instance, Pérez-Jaramillo (2019) (22) designed a greenhouse experiment with pots containing either native or agricultural soil to germinate bean seeds, either wild or domesticated. Pots without seeds served as reference bulk soils. On the other hand, Stopnisek and Shade (23) used non-adhered soil after vigorously shaking the plant roots as bulk soil.

This study observed no significant differences in bacterial diversity across bulk soils of varying physicochemical compositions and cultivation histories. Conversely, prior studies have noted a considerable impact of soil type on bacterial rhizosphere composition in wild and modern common bean accessions in Colombia and Minnesota (22, 53). This dissimilarity may be attributed to the ample contrasts in physicochemical parameters observed between the Colombian and Minnesota soils sampled, including pH (ranging from 4.7 to 8 for agricultural and native soils) and organic matter content (22, 23). Soil is often considered a critical contributor to microbial diversity in the rhizosphere of plants, with pH being a significant factor (54). In *Rhizobium legume* symbiosis, pH plays a role in selecting the microsymbiont (55). In acidic soils, *R. tropicii* species dominate the nodulation of common beans, while in soil with neutral o near neutral pH, *R. etli* and *R. phaseoli* are most common (55–57). In our experiment, slight variations in soil pH (by 0.4 units) and organic matter content may not have been significant enough to alter the bacterial communities in the bulk soils of plots A and N. Although it is well documented that the relative abundance and diversity of bacteria tend to increase as pH shifts from 4 to 8, slight pH variations are not anticipated to exert an effect (54, 58, 59). Long-term fertilization with nitrogen, phosphorus, and manure has been found to enhance the richness of the soil microbiota in alfalfa (*Medicago sativa*) but not in wheat (*Triticum*) (52). Likewise, tillage and land use could influence the soil microbiome composition; however, in the agricultural plots studied here, any possible effect of these factors on the soil microbiome was indistinguishable.

Although the dominance of the phylum Proteobacteria in the rhizosphere of Pinto Saltillo bean was unexpected, species of Proteobacteria were abundant in the rhizosphere of other plants, sharing the niche with Actinobacteria and other less represented phyla (Supplementary Figure S6). Studies based on high-throughput 16S rRNA gene sequencing have shown that additional bacterial phyla, such as Acidobacteria, Bacteroidetes, and Verrucomicrobia, are found in the rhizosphere of diverse bean cultivars (22, 60, 61).

Although taxa identified at higher taxonomic levels (Phylum, Class, Order) are more accurate and provide valuable insights into the general structure of the community, they are insufficient for describing the fundamental structural and functional units of the microbiome, such as species and their variability in terms of strains (62). However, the abundance of taxa was the result of cumulative differential abundances in the lower taxonomic categories. Multiple species or OTUs may contribute differentially to the relative abundance at the genus level; however, only a few OTUs within the genus reach high levels of representation in the community.

In the rhizosphere of the Pinto Saltillo cultivar, abundance at the genus level indicated that a group of Promoter-Growth Plant Rhizobacteria plays an important role (63). Consistently, the genera *Pseudomonas*, *Rhizobium*, *Pantoea*, and *Variovorax* were the most abundant in the rhizosphere of the cultivar Pinto Saltillo. Other genera such as *Enterobacter*, *Lellotia*, and *Stenotrophomonas*, also related to PGPR organisms, were less abundant but found in one or more rhizosphere metagenomic samples. To the best of our knowledge, these genera have also been found in different high-throughput studies but not in the microbiome of *Phaseolus vulgaris* (23). However, Rocha et al. (63) recently reported a plethora of cultivated PGPR in the rhizosphere of *Phaseolus vulgaris* L., variety “Patareco,” in agreement with the metagenomic PGPR registered in this study.

The PGPR group has been intensively studied for decades because of its beneficial effects on plant growth and development, offering opportunities for sustainable agriculture and biotechnological applications (64). It is well-known the role of strains of *Pseudomonas* in pathogen suppressing in common bean (18) and the nitrogen fixation by *Rhizobium* in symbiotic nodules (65). Other PGPRs, such as *Enterobacter*, *Pantoea*, and *Variovorax*, may produce phytohormones and antibiotics and contribute to phosphorus solubilization and root nutrient assimilation (64).

Identifying species and even strains in bacterial communities is challenging because of the limited resolution of methods based on genetic similarity or compositional profiles (62, 66). Thus, this work aimed to characterize the bacterial rhizosphere community at the species-OTU level using high-resolution shotgun metagenomic DNA sequencing and metagenome-assembled genome (MAG) reconstruction. We obtained 150 medium-to high-quality MAGs from the bulk soil, rhizosphere, and nodules distributed across the analyzed metagenomes. After eliminating redundancy, most MAGs belonged to the gamma-proteobacteria class. The *Pseudomonas* genus stands out for its exacerbated abundance in all metagenomes studied. However, obtaining *Pseudomonas* MAGs proved to be challenging. Ample variability within the *Pseudomonas* genus may hinder adequate contig binning and MAG reconstruction.

Nevertheless, MAGs corresponding to the PGPR group were obtained but functional significance of this group of MAGs was not addressed on this study. Except for *Rhizobium* MAGs, for which the nearest species was identified as *R. sophoriradicis* CCBAU 03470, its functional role in nitrogen-fixation is well known. This bacterial species was isolated from root nodules of the leguminous plant (*Sophora flavescens*) but was also found in nodules of common beans (*Phaseolus vulgaris*) in other parts of the world (67, 68). Whole genome nucleotide identity (ANIm) showed that *Rhizobium* MAGs are similar to *Rhizobium* Kim5 strains from the USA and *Rhizobium* IE4771 isolated in Mexico. These may conform to the new cosmopolitan lineage of *Rhizobium*.

Pinto Saltillo is a cultivar developed through traditional agronomic breeding techniques, selected for tolerance to rust, anthracnose, and common blight, as well as high grain yield production per hectare, estimated at 1.5 tons/ha, five years after its release (25). It is less apparent that plant hybridization genetic methods and historical domestication have been used to improve common bean cultivars by selecting their microbiome composition. This is a poorly studied area but comparing the rhizosphere microbiomes of *Fusarium*-resistant and *Fusarium*-susceptible common bean cultivars showed distinct rhizosphere microbiomes (18, 35). Additionally, rhizosphere microbiome composition was found to be dependent on the root anatomy of the rhizosphere microbiome, probably modeled by domestication practices (17). As pointed out by Fierer, “There is no “typical microbiome,” instead, there is an ample variation in the relative abundance of taxa depending on the soil and plant genotype (54). Therefore, the root microbiome of common beans is highly influenced by the agronomic features of the cultivar but is often neglected in breeding studies. Through a deeper understanding of the soil and rhizosphere communities of the Pinto Saltillo cultivar, our study provides valuable insights for future applications.

## Acknowledgements

PAPIIT-UNAM IN215908 for VG, supported this study. GLR is a doctoral student from Programa de Doctorado en Ciencias Biomédicas, Universidad Nacional Autónoma de México (UNAM), and received a fellowship (No. CVU 746216) obtained from CONAHCYT. Thanks to José Espíritu, Víctor del Moral, and Alfredo Hernández of the Unidad de Administración de Tecnologías de la Información (UATI), and Gabriela Guerrero and Luis Lozano of the Unidad de Apoyo Bioinformática (UAB), for bioinformatic assistance. Thank are extended to the undergraduate students José Luis Letechepia, Alejandro Espinoza, Juan Pablo López, and Jacob Romero for supporting soil collection, planting, and collecting bean plants at the INIFAP Experimental Field.

## Supplementary figures

**Figure S1**. (A) Location of the sampling site. Within the map of the Mexican Republic, the state of Zacatecas is highlighted, where the INIFAP Experimental Field is located (satellite photograph). A zoom is observed focusing on the sampling area, where the plots with agricultural and non-agricultural soil are indicated, within which the collection site is marked. (B) Experimental design of systematic grid sampling and location of collection sites. Bull soil and rhizosphere samples were taken at each intersection (yellow dots). Rhizosphere samples consisted of a rhizosphere pool from 9 plants. The physicochemical and soil type analysis was performed at INIFAP-Zacatecas (blue dots).

**Figure S2**. **Rarefaction curves for observed OTUs in metagenomes.** Rarefaction curves were performed with the Vegan package in R, for 17 shotgun metagenomic samples (see methods). Six soil samples sequence reads saturated the curve at 1 × 10^7^ sequences, while rhizosphere samples needed 5 × 10^7^ to reach the asymptote of the curve. This indicates that sufficient sampling sequencing depth to estimate the diversity in bulk soil and in rhizosphere.

**Figure S3. Relative abundance of bacterial phyla presents in bulk soil and rhizosphere.** Bar plots of 37 bacterial phyla. In bulk soil, the most representative phyla belong to Actinobacteria (aqua green color) and Proteobacteria (salmon color).

**Figure S4. Relative abundance of bacterial class in bulk soil and rhizosphere**. Bar graphs indicate that in the bulk soil, the predominant bacterial classes are Actinobacteria (orange color), alpha-Proteobacteria (light purple color) and beta-Proteobacteria (lime green color), while in the rhizosphere the most representative classes are gamma-Proteobacteria (green color) and alpha-Proteobacteria (light purple color) and, to a lesser extent, beta-Proteobacteria (lime green color). Bacterial classes with an abundance < 1% are represented with purple color.

**Figure S5. Absolute readings distribution of OTUs assigned to *Rhizobium*.** The absolute number of sequence readings assigned to 41 *Rhizobium* OTUs were distributed according to their abundance. *y-*axis is the log^10^ of the median sequence reads’ abundance for each OTU. *x*-axis represents the 41 OTUs assigned to *Rhizobium*.

**Figure S6**. **Relative abundance of bacterial phyla in other plants**. In the rhizosphere of other plants, the Proteobacteria (blue color) phylum is the most predominant followed by the Actinobacteria (red color) phylum.

**Figure S7. Relative abundance of bacterial classes in other plants**. The gamma-Proteobacteria class (magenta color) predominates in the rhizosphere of the common bean Pinto Saltillo. On the other hand, the alpha-Proteobacteria (king blue color), beta-Proteobacteria (sky blue color), and Actinobacteria (red color) classes are more abundant in the rhizosphere of other plants.

**Figure S8. Relative abundance of bacterial genus in other plants**. The rhizosphere of the common bean Pinto Saltillo showed a pronounced change in the relative abundance of the several bacterial genus compared to the bulk soil.

**Figure S9. MAGs quality analysis.** Box plot of the size, %GC and completeness of high-quality MAGs of the bulk soil (red color), rhizosphere (pink color) and nodules (green color). The high-quality criteria for choosing the MAGs were a size greater than 1Mb, a completeness > 60%, and a redundancy less than <10%.

## Supplementary tables

**Table S1.** Summary of the samples used for metagenome sequencing: site of origin and soil type.

**Table S2.** Physicochemical properties of agricultural and non-agricultural soils.

**Table S3.** Statistics of metagenomic sequences analysis, and distribution of read s in domains determined by Kraken2.

**Table S4**. Differential abundance of OTUs in soil and rhizosphere calculated by DESeq2.

**Table S5.** SRA accession numbers of bulk soil and rhizosphere of maize, wheat, cucumber, tomato, common bean, and *Arabidopsis*.

**Table S6**. Summary of the features of high and medium-quality MAGs.

